# Variability of pollen grains quality in oat amphiploids and their parental species

**DOI:** 10.1101/2021.07.24.453654

**Authors:** Paulina Tomaszewska, Romuald Kosina

## Abstract

The reproductive potential of oat species and their hybrid progeny (amphiploids) was evaluated during three vegetation seasons. Morphotypes and viability of pollen grains were described by means of correlations and regression, while relationships between taxa were analysed with the use of numerical taxonomy methods. Size of pollen grains varied between the growing seasons, but the relations between the taxa appeared to be stable. Viability of pollen grains was environmentally modified and showed no correlation with pollen length. In an ordination space, amphiploids were discriminated from parental species. In both group of plants, a positive correlation between the pollen size and the level of ploidy was maintained; however, along a regression line, amphiploids were located among species with a high level of ploidy and were extreme units deviating from the regression line. Developmental anomalies of pollen grains had a low frequency, with the formation of micrograins being the most common event. Such a pattern of development can prove that some pollen grains were chromosomally unbalanced. Anomalous morphotypes of pollen were more common in hybrid types than in species, including pollens with many poruses, which were found only in amphiploids. Frequencies of multiporate grains and micropollens were strongly correlated. In an ordination space, monoporate types (species) were discriminated from multiporate types (amphiploids). In general, the high level of pollen viability in amphiploids can prove their genomic stabilisation through many generations of their reproduction.

## Introduction

Research on the development and variability of pollen grains is important for understanding the mating system in plants, including their common hybrid swarms. The variability of the mating systems influences the variability and the differentiation processes observed in the populations of wild and cultivated plants (Grant 1981; Richards 1986). In cereals, the most important grasses to mankind, both natural and/or artificial selections through subsequent generations stabilise a reproduction process at the even number of chromosomes (Kushwaha et al. 2004). Such changes also apply to wild and cultivated oats as well as their artificial hybrids.

Reproductive success depends primarily on the pool of viable pollen grains available during the flowering time. This is ensured by regular meiosis, typical for fixed taxa; however, in some species, but especially in hybrids, this process can be irregular. Various anomalies related to chromosomes or activity of karyokinetic spindle were observed in meiosis of some *Avena* L. species (Sheidai et al. 2003), cultivars of *Avena sativa* L. (Baptista-Giacomelli et al. 2000a, b) or *Urochloa decumbens* (Stapf) R.D.Webster (previously classified as *Brachiaria decumbens Stapf*; Mendes-Bonato et al. 2002). These anomalies often ended with the formation of cytoplasmic separations with micronuclei and ultimately micropollens. Meiotic disturbances were more severe and more frequent in interspecific and intergeneric hybrids (Mujeeb et al. 1978; Linde-Laursen and von Bothmer 1993: Li et al. 2005); however, their pattern was similar. Thus, studies on the frequency of micronuclei and micropollens can be a good tool for evaluation of the reproductive quality of the plant. Meiotic disorders can also be found even in one anther sac, where the microsporocyte mother tissue forms a mosaic of cells with different numbers of chromosomes. Such an event was found in the amphiploid *Avena abyssinica* × *A. strigosa* (Thomas and Peregrine 1964) and in *Allium senescens* subsp. *montanum* (Pohl) Holub (synonym of accepted *Allium senescens* L.) (Malecka 2008).

In the highly polyploid species or hybrids, including amphiploids with a doubled chromosome set, a change from generative into apomictic reproduction was described (Grant 1981; Quarin et al. 2001; Ma et al. 2009). A very special morphogenesis of pollen grains in the form of multiporate units has been interpreted as related to apomixis (Ma et al. 2009). Multiporate grains appeared to be ineffective due to the germination of pollen tubes through several pores and their mutual competition under a tube growth (Florek 2013; Kosina et al. 2014). A pattern of porus formation is probably determined by metabolism and deposition of callose before a porus morphogenesis (Teng et al. 2005). It was also evidenced that the callose deposition in a pollen grain varied among grasses (Warzych 2001; Kłyk 2005).

Evaluation of the reproductive stability of plant hybrids and their relatives is important for breeding purposes. Therefore, this article presents comparative analyses of pollen quality in oat species showing a broad ploidy spectrum and their amphiploids. Some components of the mating system of oat species and amphiploids were assessed taking into account the morphology and viability of pollen grains as indicators of hybrid vigor.

## Materials and methods

Oat amphiploids and their parental species, listed in Table 1, were cultivated on small plots in the grass collection (Wrocław, SW Poland) maintained by R. Kosina. During the plot experiments, the plants were grown under the same soil-climatic conditions. Mature stamens were sampled from plants at the turn of June and July, for the three consecutive growing seasons. Each year, mature stamens taken from 5 plants of each of the 15 accessions (OTUs, Operational Taxonomic Units) were analysed. Stamens were always collected from the lower flower of spikelet. Pollen grains isolated from anther were placed on microscope slides, stained with 0,5% v/v acetocarmine or 0,01% v/v acridine orange, mounted in glycerin, and covered with a coverslip. The botanical nomenclature of studied species was applied according to https://npgsweb.arsgrin.gov/gringlobal/taxon/taxonomysearch.aspx, accessed on 21 July 2021, and http://www.theplantlist.org/, accessed on 21 July 2021:

*A. fatua* L.
*A. sterilis* L.
*A. sativa* L.
*A. barbata* Pott ex Link
*A. abyssinica* Hochst.
*A. magna* H.C. Murphy & Terrell
*A. strigosa* Schreb.
*A. longiglumis* Durieu
*A. eriantha* Durieu

**Table 1.** List of studied accessions of oat amphiploids and their parental species

| Amphiploid and parental species | Abbreviation | Accession number | Origin | Ploidy level |
| --- | --- | --- | --- | --- |
| *A. barbata* × *A. sativa* subsp. *nuda* | *b/sn* | CIav7903** | - | octoploid |
| *A. eriantha* × *A. sativa* | *e/sa* | PI458781** | - | octoploid |
| *A. barbata* × *A. sativa* | *b/sa* | CIav7901** | - | hexaploid |
| *A. fatua* × *A. sterilis* | *f/ste* | CIav9367** | - | hexaploid |
| *A. magna* × *A. longiglumis* | *m/l* | CIav9364** | - | hexaploid |
| *A. abyssinica* × *A. strigosa* | *a/str* | CIav7423** | - | tetraploid |
| *A. fatua* | *Af* | RK | Poland | hexaploid |
| *A. sterilis* | *Aste* | PI311689** | Israel | hexaploid |
| *A. sativa* | *Asa* | RK | Poland | hexaploid |
| *A. barbata* | *Ab* | AVE1938* | Spain | tetraploid |
| *A. abyssinica* | *Aa* | PI331373** | Ethiopia | tetraploid |
| *A. magna* | *Am* | 1786*** | Morocco | tetraploid |
| *A. strigosa* | *Astr* | 51624* | Belgium | diploid |
| *A. longiglumis* | *Al* | PI367389** | Portugal | diploid |
| *A. eriantha* | *Ae* | CIav9051** | England | diploid |

\* Bundesanstalt für Züchtungsforschung an Kulturpflanzen, Braunschweig, Germany; \*\* National Small Grains Collection, Aberdeen, Idaho, USA; \*\*\* Vavilov Institute of Plant Industry, St. Petersburg, Russia; RK – collected by R. Kosina

### Microscopy

The slides were examined under an Olympus BX-60 epifluorescence microscope (Hamburg, Germany), using both white and UV light. Images were taken with an Olympus E-520 camera (Olympus Imaging Europe, Hamburg, Germany).

### Statistical analysis

OTUs have been described by the following characteristics of pollen grains:

- viability (all viable, dysfunctional and dead pollen grains taken from anther were counted),
- length of the pollen grain (measured for forty randomly selected pollen grains from anther),
- impaired pollen grains (all micrograins, unreduced pollen grains, and grains of irregular shape were counted),
- number of poruses in pollen grains.

The OTUs were arranged into an ordination space with application of non-metric multidimensional scaling (Kruskal 1964; Rohlf 1981) and according to Rohlf ’s numerical approach (Rohlf 1994). The correlation and regression analyses were made according to KWIKSTAT 4 procedure (Elliot 1994).

## Results

### Viability and size of pollen grains

The viability of the pollen grains was determined using the acetocarmine staining technique, which allowed the grains to be classified into three groups:

- normal, viable pollen - stained red, completely filled with cytoplasm,
- dysfunctional, defective pollen - weakly coloured, partially filled with cytoplasm,
- empty, dead pollen - uncoloured, with no cytoplasm, usually wrinkled or cracked.

Defective and dead pollen grains were treated together as non-viable pollens in statistical analyses.

Table 2 presents the results of analyses concerning the percentage of viable pollen grains in the examined accessions, taking into account the three consecutive growing seasons. Accessions of both amphiploids and their parental species were characterised by a high level of pollen grain viability. The average percentage of viable pollen for three years was over 90% in all examined accessions. Amphiploids showed a slightly lower viability of pollen grains compared to their parental species. The percentage of viable pollen in species ranged from 96.3 to 98.9%, and in amphiploids the range was 90.5 - 96.0%. Octoploid amphiploids were characterised by the lowest pollen viability: the average percentage of pollen completely filled with cytoplasm in *b/sn* and *e/sa* was 91.1% and 90.5%, respectively. In some growing seasons, some species and amphiploids showed pollen grain viability below 90% or pollen was not obtained for analysis due to problems with seed germination or premature drying of plants due to the drought. Over the three-year period, the range of variation of pollen viability between the three growing seasons was greatest in the amphiploid *f/ste* (from 82.7 to 98.0%) and the hexaploid species *Asa* (from 88.5 to 99.0%).

**Table 2.**
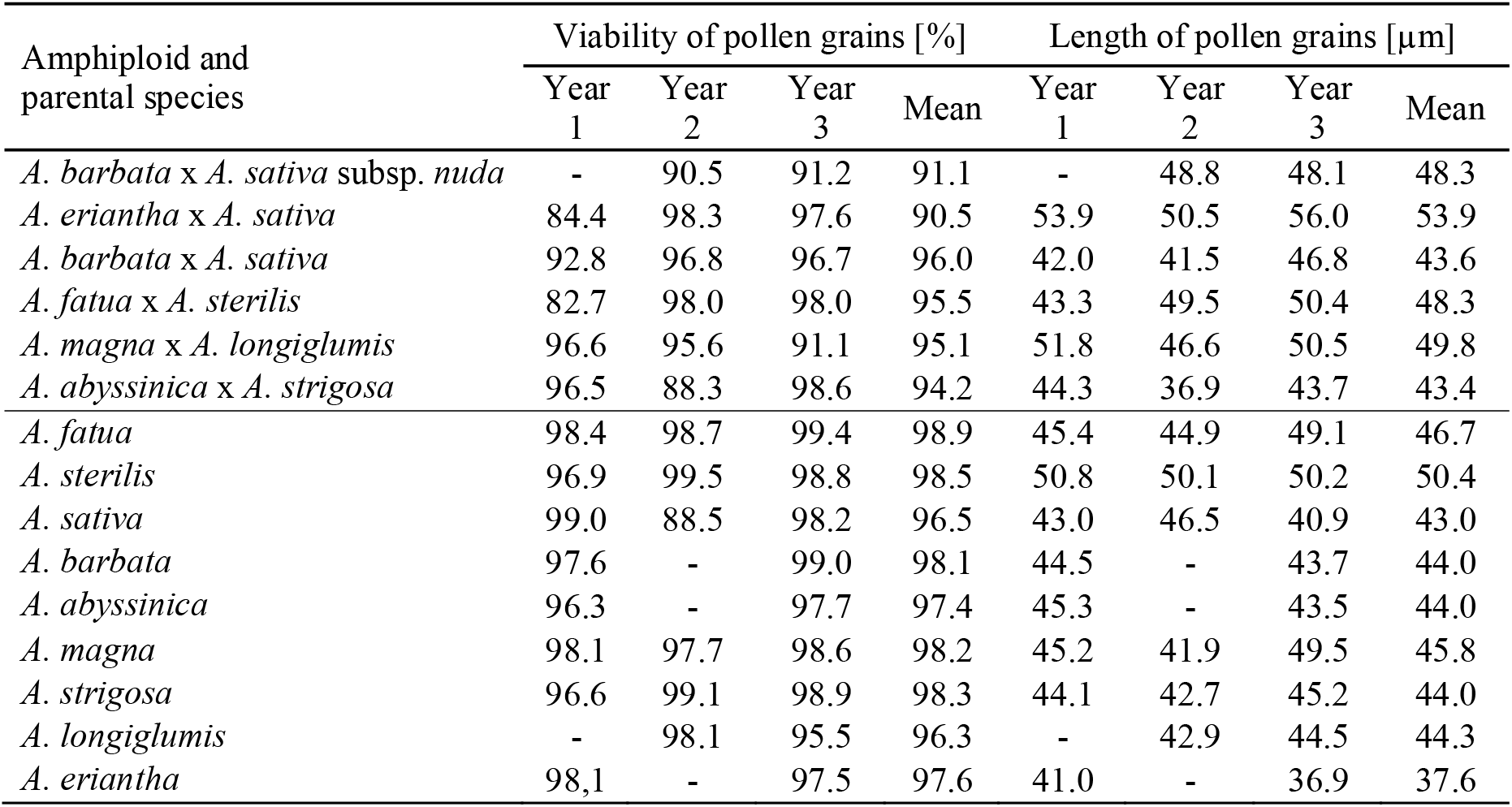
Viability and size of pollen grains of oat amphiploids and their parental species

| Amphiploid and parental species | Viability of pollen grains [%] |  |  |  | Length of pollen grains [µm] |  |  |  |
| --- | --- | --- | --- | --- | --- | --- | --- | --- |
|  | Year 1 | Year 2 | Year 3 | Mean | Year 1 | Year 2 | Year 3 | Mean |
| *A. barbata* x *A. sativa* subsp. *nuda* | - | 90.5 | 91.2 | 91.1 | - | 48.8 | 48.1 | 48.3 |
| *A. eriantha* x *A. sativa* | 84.4 | 98.3 | 97.6 | 90.5 | 53.9 | 50.5 | 56.0 | 53.9 |
| *A. barbata* x *A. sativa* | 92.8 | 96.8 | 96.7 | 96.0 | 42.0 | 41.5 | 46.8 | 43.6 |
| *A. fatua* x *A. sterilis* | 82.7 | 98.0 | 98.0 | 95.5 | 43.3 | 49.5 | 50.4 | 48.3 |
| *A. magna* x *A. longiglumis* | 96.6 | 95.6 | 91.1 | 95.1 | 51.8 | 46.6 | 50.5 | 49.8 |
| *A. abyssinica* x *A. strigosa* | 96.5 | 88.3 | 98.6 | 94.2 | 44.3 | 36.9 | 43.7 | 43.4 |
| *A. fatua* | 98.4 | 98.7 | 99.4 | 98.9 | 45.4 | 44.9 | 49.1 | 46.7 |
| *A. sterilis* | 96.9 | 99.5 | 98.8 | 98.5 | 50.8 | 50.1 | 50.2 | 50.4 |
| *A. sativa* | 99.0 | 88.5 | 98.2 | 96.5 | 43.0 | 46.5 | 40.9 | 43.0 |
| *A. barbata* | 97.6 | - | 99.0 | 98.1 | 44.5 | - | 43.7 | 44.0 |
| *A. abyssinica* | 96.3 | - | 97.7 | 97.4 | 45.3 | - | 43.5 | 44.0 |
| *A. magna* | 98.1 | 97.7 | 98.6 | 98.2 | 45.2 | 41.9 | 49.5 | 45.8 |
| *A. strigosa* | 96.6 | 99.1 | 98.9 | 98.3 | 44.1 | 42.7 | 45.2 | 44.0 |
| *A. longiglumis* | - | 98.1 | 95.5 | 96.3 | - | 42.9 | 44.5 | 44.3 |
| *A. eriantha* | 98.1 | - | 97.5 | 97.6 | 41.0 | - | 36.9 | 37.6 |

The amphiploids and their parental species varied in the pollen size. Oat pollen grains were usually slightly elongated, elliptical in shape. Thus, the measurements of their length were sufficient to describe size of pollen grains, and these data are included in Table 2. The average length of grains ranged from 37.6 to 53.9 μm. The smallest grains were observed in *Ae,* while the largest pollen was found in the octoploid *e/sa.* The mean length of the elliptical grains in the amphiploid *e/sa* was over 50 μm in all years. The size of pollen grains varied not only between the analysed accessions, but also between years. The latter depended on the weather conditions, such as temperature and water supply, during the development of pollen grains.

The Pearson’s correlation coefficients matrix of the pollen grain characteristics (pollen viability and their average length in particular growing seasons) is presented in Table 3. The average length of pollen grains in three consecutive growing seasons were correlated with each other. The significant correlations of the pollen grain size between different growing seasons mean that this feature was a stable characteristic of the analysed taxa. The lack of correlation between the viability of pollen grains in individual years indicated a random, non-directional, high environmental variation of this feature. The matrix of the Pearson’s correlation coefficients for pollen grain characteristics was used to distribute the fifteen studied oat accessions (OTUs) in an ordination space along three axes (***x***, ***y***, ***z***) using the Kruskal’s non-metric multidimensional scaling (nmMDS). The diagram of the minimum spanning tree (MST) (Fig. 1) shows the location of OTUs connected to each other by the smallest links. The smallest distance was observed between the OTUs *Aa* and *Ab.* The location of *Asa* in the vicinity of the taxon with small pollen grains is probably conditioned by the inter-variety variability of this cultivated species. The extreme position of the amphiploid *e/sa* was mainly determined by its largest pollen grains. In the diagram, a large distance between the two amphiploids *e/sa* and *f/ste,* which showed similar viability of pollen grains in different growing seasons, was due to the different size of pollen grains in both taxa. The diagram also shows a certain directional difference between species and their progeny - the species formed a group located in the center of the diagram, while amphiploids were scattered outside (Fig. 1). This means that a pattern of seasonal variation of two analysed characteristics varied between oat species and amphiploids; however, this was not complete discrimination as *Aa* and *Ae* appeared to be outside of the center.

**Table 3.** Pearson’s correlation coefficients matrix of the two pollen grain characteristics, viability and length, in the three growing seasons

|  | V1 | V2 | V3 | L1 | L2 |
| --- | --- | --- | --- | --- | --- |
| V2 | -0.24 |  |  |  |  |
| V3 | -0.01 | 0.25 |  |  |  |
| L1 | -0.26 | 0.31 | -0.32 |  |  |
| L2 | -0.46 | 0.27 | -0.28 | 0.60* |  |
| L3 | -0.58* | 0.57* | -0.11 | 0.73*** | 0.55* |

V1, V2, V3 - viability of pollen grains in the three growing seasons; L1, L2, L3 - length of pollen grains in the three growing seasons; \*, \*\*\* - significance levels at $\alpha = 0.05, 0.001$ , respectively

**Fig. 1.**
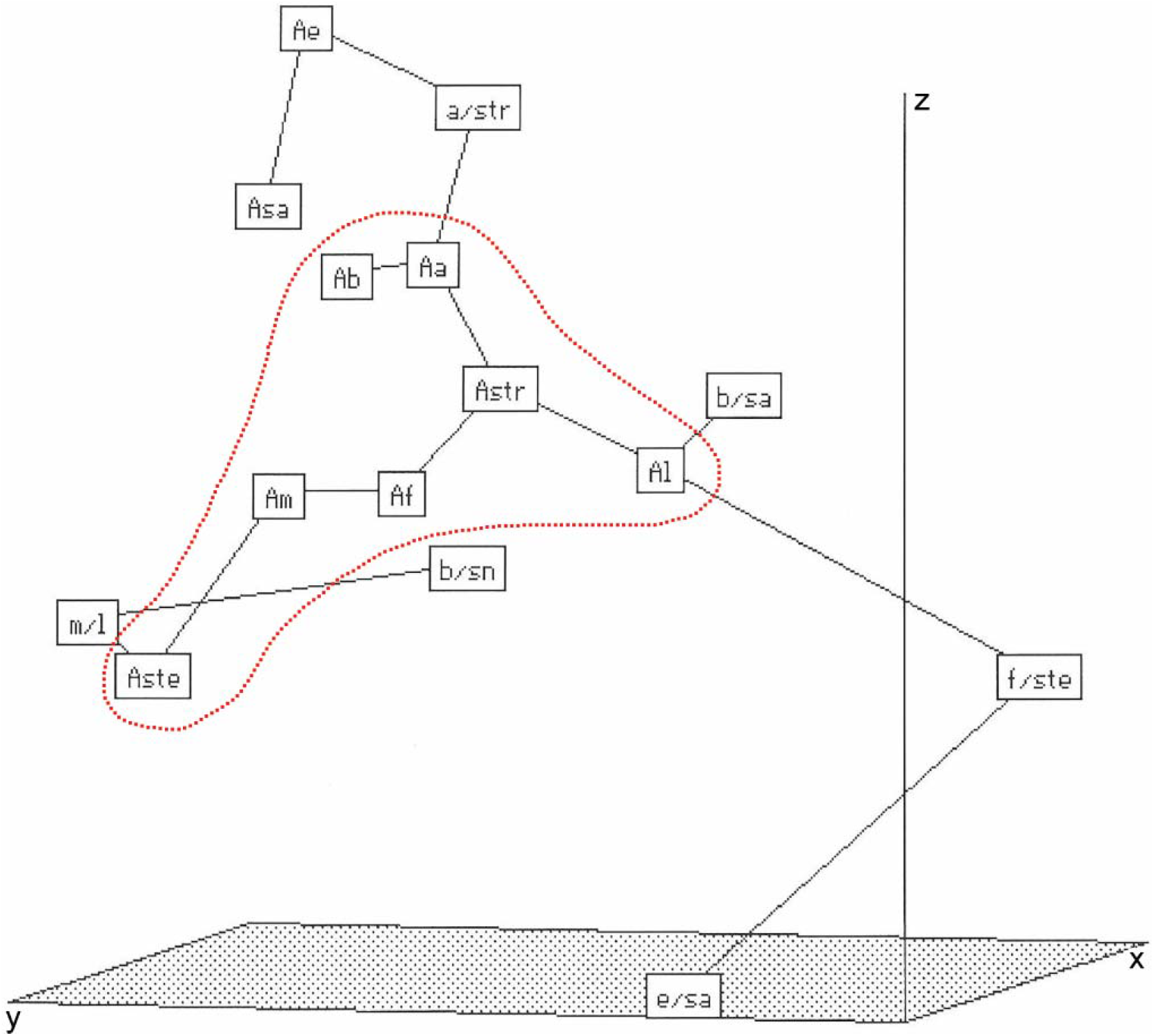
Minimum spanning tree (MST) of amphiploids and parental species (OTUs) of the genus *Avena* in an ordination space (*x*, *y*, and *z-*axes) and created by application of Kruskal’s non-metric multidimensional scaling method (Rohlf 1994). OTUs were described by pollen viability and pollen length in the three growing seasons. Abbreviations in the Table 1. The distances between pairs of OTUs in the minimum spanning tree (MST) are as follows (extreme distances are bolded): *b/sn* - *m/l* 0.869; *m/l* - *Aste* 0.775; *Aste* - *Am* 0.703; *Am* - *Af* 0.245; *Af* - *Astr* 0.398; *Astr* - *Aa* 0.469; ***Aa* - *Ab* 0.205**; *Astr* - *Al* 0.489; *Al* - *b/sa* 0.296; *Aa* - *a/str* 0.602; *a/str* - *Ae* 0.664; *Ae* - *Asa* 0.743; *Al* - *f/ste* 1.040; ***f/ste* - *e/sa* 1.231**

### Regression of pollen length versus ploidy

The research showed a significant correlation between the length of pollen grains and the level of ploidy provided for the root tissues (Table 1; P. Tomaszewska pers. comm.).The value of the Pearson’s correlation coefficient for a random sample of *n*=15 OTUs was *r*=0.78, which indicates the significant regression relationship between the variables (Fig. 2). Low values of both characteristics were observed in di- and tetraploid species, while hexaploid species and amphiploids were located in the right part of the diagram showing higher values. In addition, for the extreme OTUs, *e/sa* and *b/sn,* a high frequency of multiporate pollen grains was noted (see Tables 5 and 6).

**Fig. 2.**
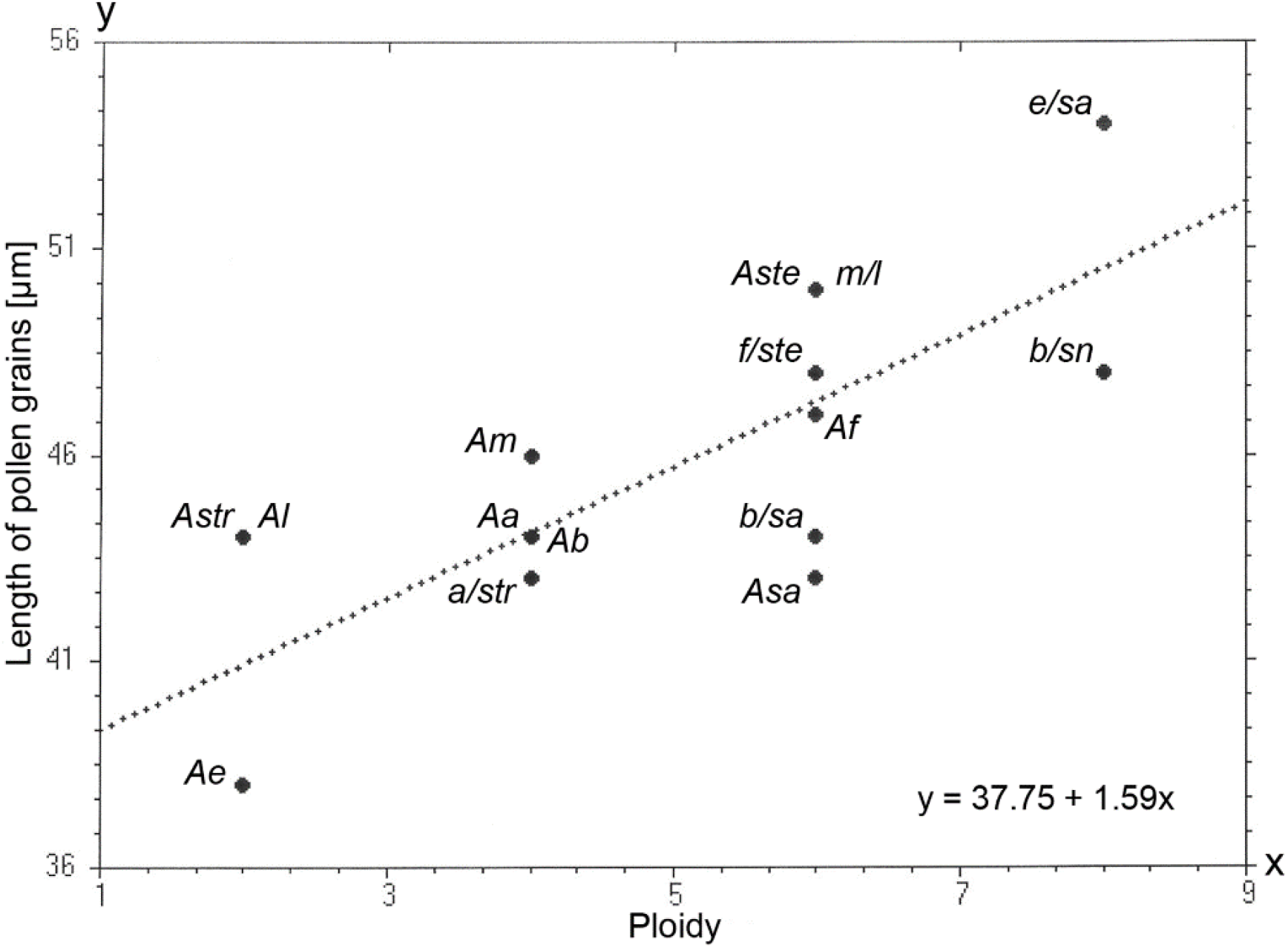
A diagram of the linear regression showing relationship between pollen grain length and ploidy level of oat amphiploids and parental species. Length of pollen grains in μm

### Anomalous pollen grains

Apart from normal pollen grains, some grains showed developmental disorders, and their examples are presented in Fig. 3. In all the examined accessions, pollen micrograins were observed (Fig. 3a), but their percentage share in the entire pool of pollen grains tested did not exceed 1% (Table 4). *Astr,* as well as three amphiploids (*e/sa, m/l* and *a/str*), showed very large, unreduced pollen grains. Some such grains are shown in Fig. 3b,d,e,f. Their considerable size may prove a higher level of ploidy of pollen stem cell or non-reduction of gametes. It may also be caused by disturbances in the formation of cell walls during subsequent cytokineses in the microsporogenesis process, which was confirmed by the irregular shape and depressions in the surface of some large pollen grains (Fig. 3c,g,h). However, the frequency of large pollen grains was low and did not exceed 1%. In some cases, the irregular shape of the pollen grains suggests an asymmetric division that led to the formation of micropollens (Fig. 3j). In the amphiploid *e/sa,* a single case of a huge, highly elongated pollen was recorded (Fig. 3k). It was estimated to be about 4-5 times larger than a normally developed grain, and had seven poruses, including two connected. Anomalous pollen grains were observed in all amphiploids, but also in one hexaploid and three tetraploid species (Table 4). However, they constituted a small percentage of the total pollen pool analysed. The highest amount of pollen of irregular shape was observed in octoploids, but their frequency was only 0.06% and 0.05% in *e/sa* and *b/sn,* respectively. Micrograins appeared to be most common among other anomalies. Their average frequency was in amphiploids 0.51%, while in the parental species it was distinctly lower 0.21%.

**Fig. 3.**
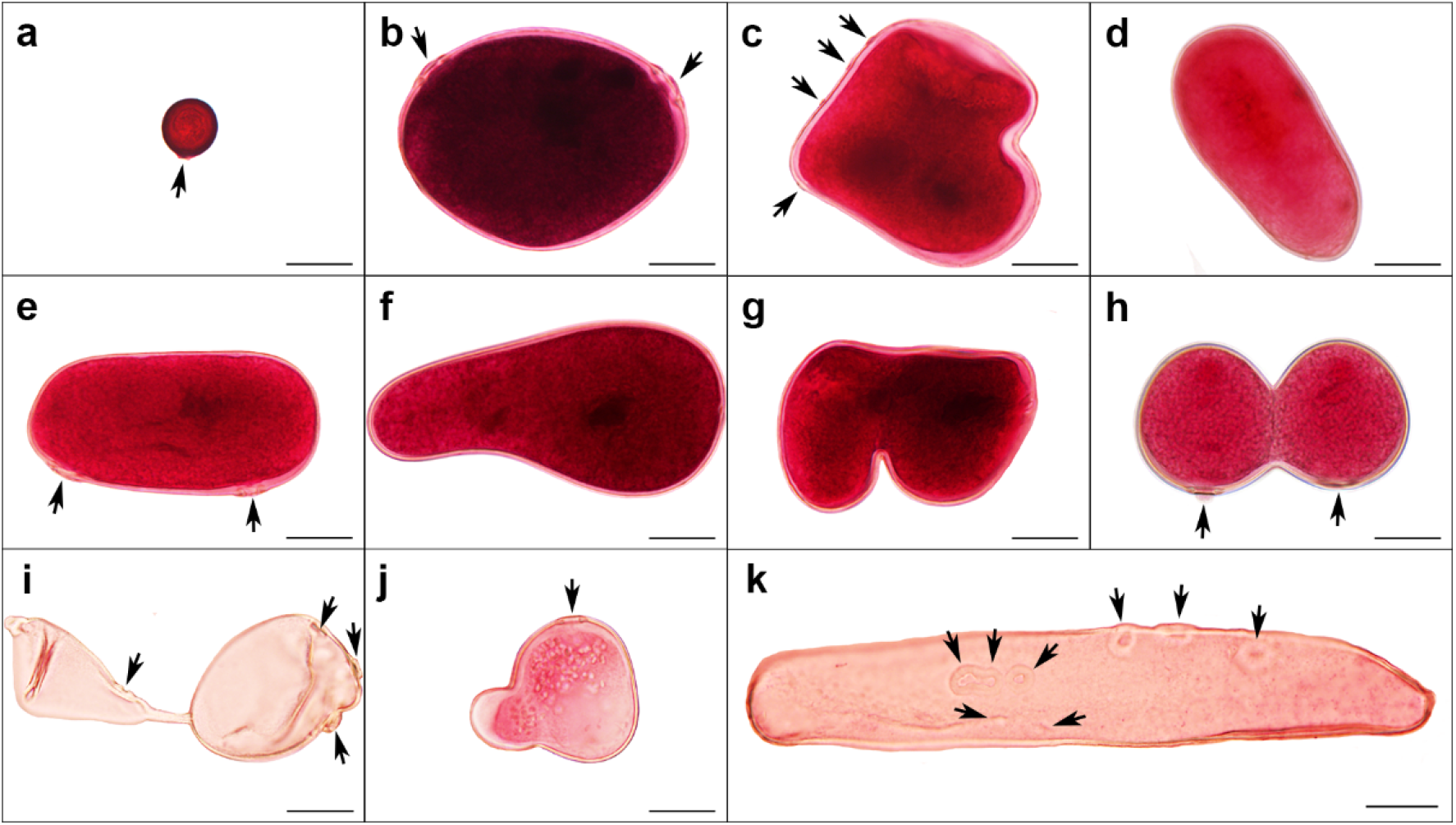
Anomalous morphotypes of pollen grains. a – *A. fatua* × *A. sterilis;* b,c – *A. eriantha* × *A. sativa;* d - *A. abyssinica* × *A. strigosa;* e - *A. eriantha* × *A. sativa;* f,g – *A. eriantha* × *A. sativa;* h - *A. abyssinica* × *A. strigosa;* i - *A. barbata* × *A. sativa;* j – *A. fatua* × *A. sterilis;* k - *A. eriantha* × *A. sativa.* Acetocarmine staining, black arrows indicate poruses. Scale bars = 20μm

**Table 4.** Frequency of anomalous pollen grains (three-year average)

| Amphiploids and parental species | Anomalous pollen grains [%] |  |  |
| --- | --- | --- | --- |
|  | micropollens | unreduced | irregular |
| *A. barbata* × *A. sativa* subsp. *nuda* | 0.89 | 0 | 0.05 |
| *A. eriantha* × *A. sativa* | 0.62 | 0.02 | 0.06 |
| *A. barbata* × *A. sativa* | 0.28 | 0 | 0.01 |
| *A. fatua* × *A. sterilis* | 0.90 | 0 | 0.01 |
| *A. magna* × *A. longiglumis* | 0.12 | 0.01 | 0.02 |
| *A. abyssinica* × *A. strigosa* | 0.22 | 0.02 | 0.01 |
| *A. fatua* | 0.18 | 0 | 0.01 |
| *A. sterilis* | 0.30 | 0 | 0 |
| *A. sativa* | 0.19 | 0 | 0 |
| *A. barbata* | 0.03 | 0 | 0.02 |
| *A. abyssinica* | 0.46 | 0 | 0.03 |
| *A. magna* | 0.17 | 0 | 0.002 |
| *A. strigosa* | 0.24 | 0.01 | 0 |
| *A. longiglumis* | 0.09 | 0 | 0 |
| *A. eriantha* | 0.23 | 0 | 0 |

### Number of poruses in cell walls of pollen grains

As shown in Fig. 3, irregularly shaped pollen grains often had more than one porus. The analysis of the number of poruses was performed under UV light after staining the pollen with acridine orange. Normal pollen grains of the analysed oat accessions had one porus (Fig. 4a). However, in all tested amphiploids, also in their normal pollen grains, two poruses were observed sporadically. This phenomenon concerned only amphiploids and was absent in the parental species. The position of the poruses in relation to each other was different. Sometimes they were distant (Fig. 4c) or close to each other (Fig. 4e), and in some cases connected to each other (Fig. 4b,d). The diameters of both poruses were similar or different (Fig. 4c,f). In the amphiploid *b/sn,* single cases of pollen with three poruses (Fig. 4f) and four poruses were reported. Amphiploid *e/sa* also had pollen grains with a greater number of poruses. An extreme case of the multiporate grain was noted in this amphiploid (Fig. 3k). Pollen with three and four poruses were not observed in the remaining four amphiploids. Table 5 shows that the frequency of pollen grains with an increased number of poruses was low. *b/sn* showed the highest frequency of pollen grains with many poruses in comparison to other amphiploids.

**Fig. 4.**
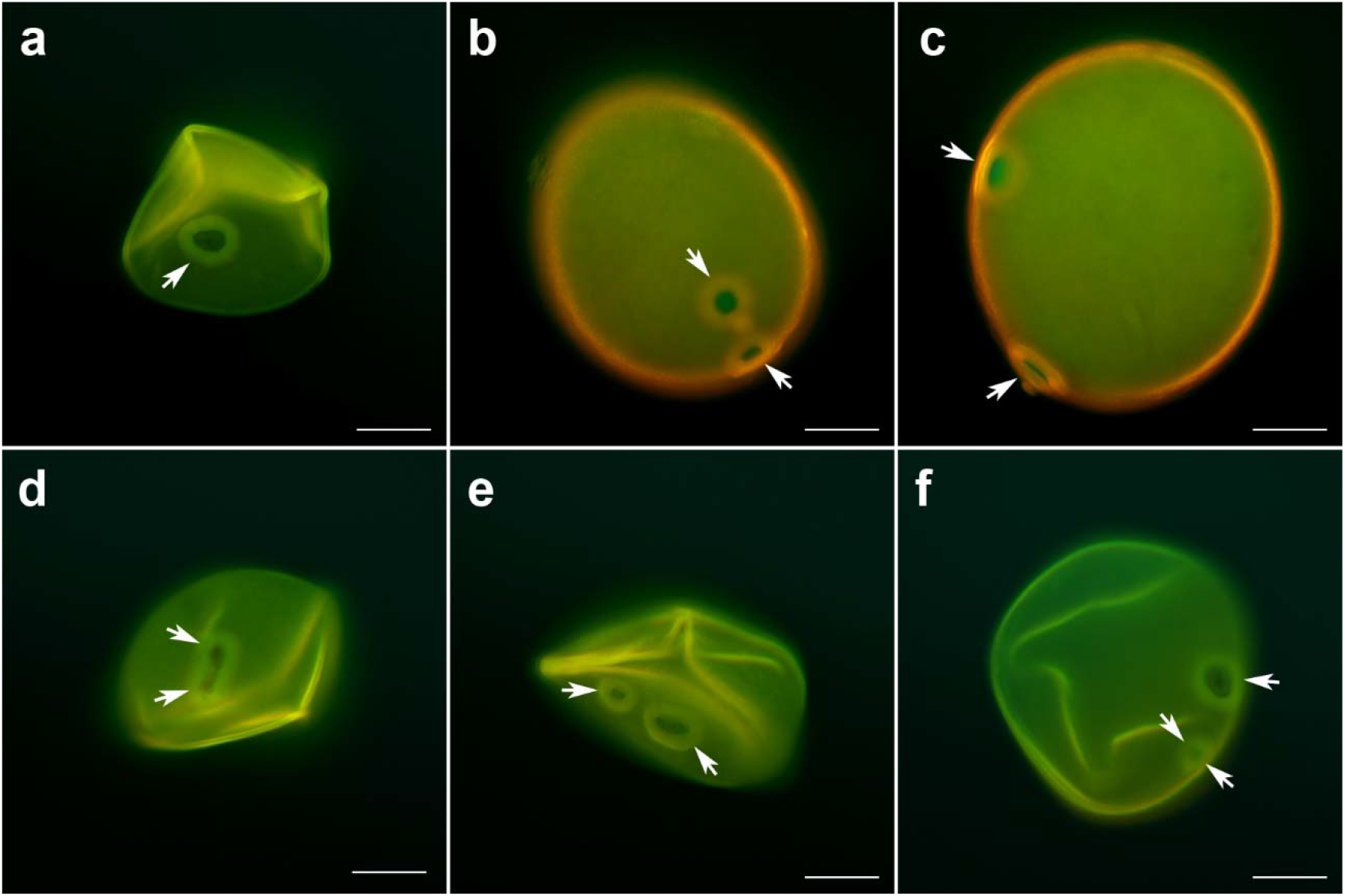
Different morphotypes, number and position of poruses in pollen of oat amphiploids. a - *A. magna* × *A. longiglumis,* pollen with one porus; b,c - *A. eriantha* × *A. sativa,* pollens with two poruses; d - *A. magna* × *A. longiglumis,* pollen with two poruses; e - *A. magna* × *A. longiglumis,* pollen with two poruses; f - *A. barbata* × *A. sativa* subsp. *nuda,* pollen with three poruses. Acridine orange fluorescence. White arrows indicate poruses. Scale bars = 20μm

**Table 5.** Frequency of pollen grains with a certain number of poruses (%)

| Amphiploids and<br>parental species | 1 porus | 2 poruses | 3 poruses | 4 poruses | > 4 poruses |
| --- | --- | --- | --- | --- | --- |
| *A. barbata* × *A. sativa* subsp. *nuda* | 97.1 | 2.28 | 0.54 | 0.13 | 0 |
| *A. eriantha* × *A. sativa* | 99.4 | 0.43 | 0.11 | 0 | 0.05 |
| *A. barbata* × *A. sativa* | 99.9 | 0.11 | 0 | 0 | 0 |
| *A. fatua* × *A. sterilis* | 99.4 | 0.62 | 0 | 0 | 0 |
| *A. magna* × *A. longiglumis* | 99.8 | 0.22 | 0 | 0 | 0 |
| *A. abyssinica* × *A. strigosa* | 99.8 | 0.21 | 0 | 0 | 0 |
| *A. fatua* | 100 | 0 | 0 | 0 | 0 |
| *A. sterilis* | 100 | 0 | 0 | 0 | 0 |
| *A. sativa* | 100 | 0 | 0 | 0 | 0 |
| *A. barbata* | 100 | 0 | 0 | 0 | 0 |
| *A. abyssinica* | 100 | 0 | 0 | 0 | 0 |
| *A. magna* | 100 | 0 | 0 | 0 | 0 |
| *A. strigosa* | 100 | 0 | 0 | 0 | 0 |
| *A. longiglumis* | 100 | 0 | 0 | 0 | 0 |
| *A. eriantha* | 100 | 0 | 0 | 0 | 0 |

### Correlation between pollen grain characteristics

For all the pollen characteristics (arithmetic averages from three growing seasons), the Pearson’s correlation coefficient matrix was calculated. Table 6 shows that the viability of pollen grains was positively correlated only with pollen grains having one porus. There were also negative correlation coefficients between pollen viability and all other characteristics. This means that taxa which produce micropollens, non-reduced pollen grains, irregular pollen and multiporate grains will show a lower ability to germinate pollen grains. Disturbances in pollen development significantly affect its viability. The viability of the pollen was also negatively correlated with the length of the pollen grains. In plants with a higher level of ploidy, and thus having larger pollen grains, a reduced pollen viability can be observed comparing to diploid species producing haploid pollen grains. The presence of micrograins was positively correlated with irregular, three- and four-porus pollen, and negatively correlated with one-porus pollen. Moreover, a positive correlation was demonstrated between the presence of pollen with more than four poruses and the length of pollen grains and non-reduced pollen. Other positive correlations occurred between irregular pollen and the length of the pollen grains, as well as between multiporate pollen grains. On the other hand, the presence of irregular pollen was negatively correlated with one-porus grain. When analysing the relationships between the characteristics of pollen grains related to the number of poruses, a negative correlation was noted between one-porus pollen and pollen with two, three and four poruses. The development of two-, three- and four-porus grains was positively correlated. All these data clearly show that the larger the pollen, and thus the higher the level of ploidy, the more disturbances in microsporogenesis, leading to the formation of anomalous pollen grains.

**Table 6.** Pearson’s correlation coefficient matrix for the ten characteristics of pollen grains (three growing seasons)

|  | V | L | MP | UR | IR | 1P | 2P | 3P | 4P |
| --- | --- | --- | --- | --- | --- | --- | --- | --- | --- |
| L | -0.51* |  |  |  |  |  |  |  |  |
| MP | -0.66** | 0.42 |  |  |  |  |  |  |  |
| UR | -0.52* | 0.33 | 0.03 |  |  |  |  |  |  |
| IR | -0.78*** | 0.58* | 0.59* | 0.37 |  |  |  |  |  |
| 1P | 0.73*** | -0.31 | 0.73*** | 0.01 | -0.64** |  |  |  |  |
| 2P | -0.73*** | 0.32 | 0.75*** | -0.01 | 0.62** | -1.00*** |  |  |  |
| 3P | -0.67** | 0.28 | 0.63** | -0.03 | 0.65** | -0.98*** | 0.96*** |  |  |
| 4P | -0.56* | 0.16 | 0.57* | -0.15 | 0.52* | -0.96*** | 0.95*** | 0.98*** |  |
| >4P | -0.56* | 0.59* | 0.30 | 0.60* | 0.67** | -0.10 | 0.08 | 0.13 | -0.07 |

V – viability of pollen grains, L – length of pollen grains, MP - micropollens, UR – unreduced pollen grains, IR – irregular pollen grains, 1P – pollens with one porus, 2P – pollens with two poruses, 3P – pollens with three poruses, 4P – pollens with four poruses, > 4P – pollens with more than four poruses

The Pearson’s correlation coefficient matrix of ten pollen grain characteristics (Table 6) was used to distribute the fifteen studied oat accessions (OTUs) in the space of three ordination axes using the Kruskal’s non-metric multidimensional scaling (nmMDS). A diagram of the minimum spanning tree (MST) (Fig. 5) shows the location of the OTUs. The nmMDS analysis revealed significant variability between the taxa studied here. Diploid *Ae* and two octoploids *b/sn* and *e/sa* occupy the opposite positions in the MST diagrams. The position of *Ae* in the diagram was determined by the very small size of pollen grains, as well as the complete absence of non-reduced and irregular pollen grains in the analysed pollen pool from three growing seasons. The other two diploids, *Al* and *Astr,* also showed no significant disturbances in the development of pollen grains, but it was the size of the pollen grains that influenced the placement of the OTUs in diagram. *Astr* was located close to the tetraploid species *Am* and *Aa.* In turn, *Al* showed similarity in the values of the studied traits to the amphiploid *b/sa.* The phenotypic distance between these two OTUs was very small. The second extreme group in MST diagrams was created by octoploids. Their location in the ordination space was due to the slightly reduced viability of pollen grains, as well as the higher frequency of irregular pollen grains compared to the other studied accessions. Moreover, both extreme amphiploids (*b/sn, e/sa*) were distinguished from other taxa by the higher frequency of multiporate pollens. Despite this, there was a clear discrimination between both OTUs. It may result from significant differences in the size of pollen grains between the accessions of the studied species and amphiploids. The length of the pollen grains was greatest in the amphiploid *e/sa.* In turn, *b/sn,* despite the same level of ploidy as *e/sa,* produced pollen of a size comparable to hexaploid amphiploids. Such a distribution of both OTUs could also be influenced by the differences in the percentage share of micropollens and non-reduced pollen grains in the studied pools. The small distance between *Ab* and *Astr* indicates their phenotypic similarity of the analysed traits. The viability of pollen grains did not differentiate them from other studied accessions, but both species were characterised by the production of pollen of a similar size. It is clear that mono- and multiporate taxa (species versus hybrid units) were well discriminated in the diagram.

**Fig. 5.**
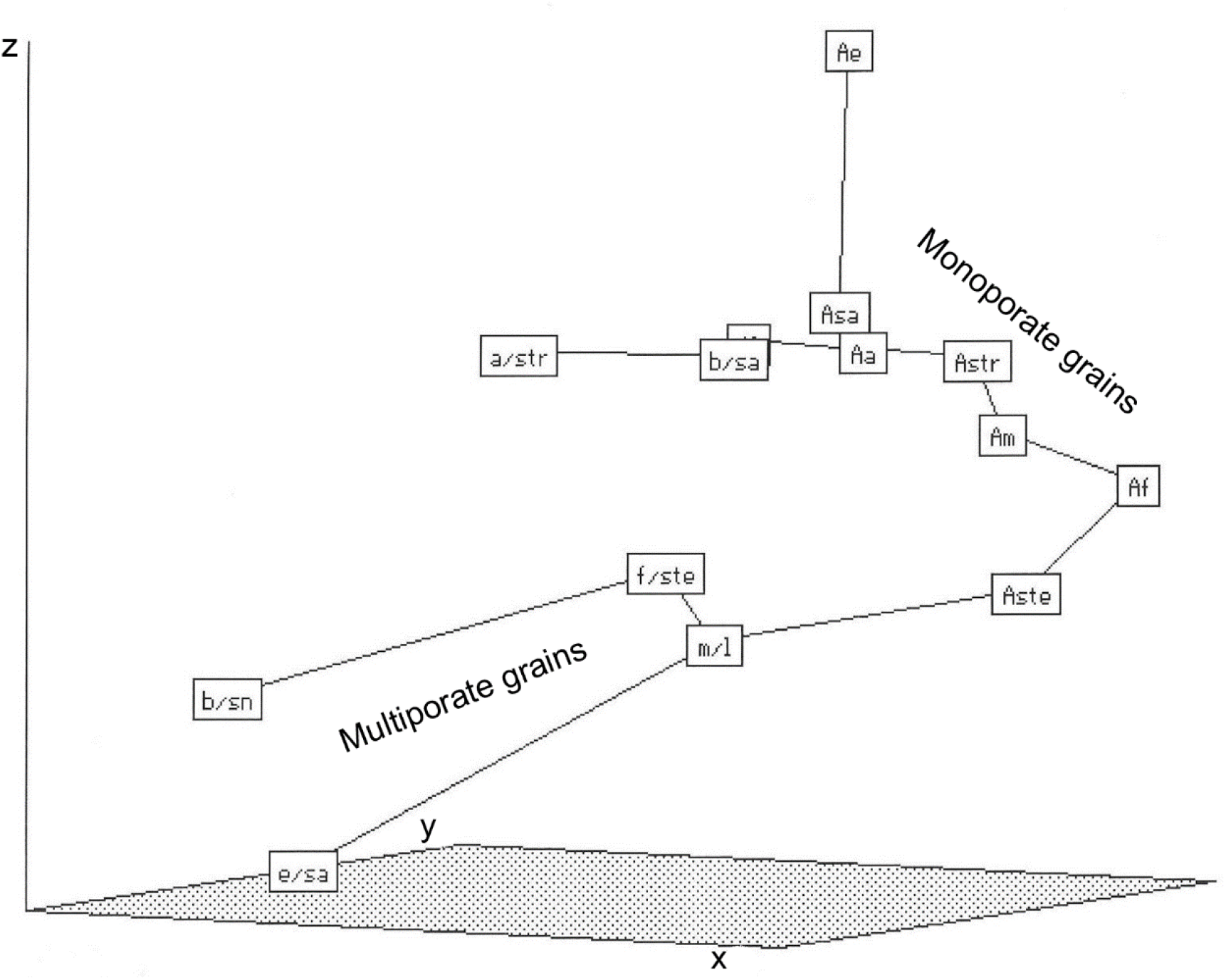
Minimum spanning tree (MST) of amphiploids and parental species (OTUs) of the genus *Avena* in an ordination space (*x*-, *y-,* and *z*-axes) and created by application of Kruskal’s non-metric multidimensional scaling method (Rohlf 1994). OTUs were described by ten traits of pollen (viability, length, micropollens, unreduced pollen, irregular pollen, pollen with 1 porus, pollen with 2 poruses, pollen with 3 poruses, pollen with 4 poruses, pollen with more than 4 poruses). Abbreviations in the Table 1. Some OTUs are hidden, *Ab* behind *Astr* and *Al* behind *b/sa.* The distances between pairs of OTUs in the minimum spanning tree (MST) are as follows (extreme distances are bolded): *b/sn* - *f/ste* 1.088; *f/ste* - *m/l* 0.460; *m/l* - *Aste* 0.639; *Aste* - *Af* 0.722; *Af* - *Am* 0.331; *Am* - *Ab* 0.463; ***Ab* - *Astr* 0.002**; *Astr* - *Aa* 0.231; *Aa* - *Al* 0.233; *Al* - *b/sa* 0.096, *Aa* - *Asa* 0.236; *b/sa* - *a/str* 0.505; *Asa* - *Ae* 1.112; ***m/l* - *e/sa* 1.265**

## Discussion

The pollen variability (Dzyuba et. al. 2006; Karabournioti et. al. 2007; Costa et al. 2016) can be evaluated by measuring pollen viability, assessing its morphology, and studying pollen-stigma compatibility reactions, thus obtaining important information on the plant breeding systems (Kosina and Tomaszewska 2015). Other biological parameters, such as pollen size (De Storme et al. 2013) and number of poruses (Kihara 1982; Kalinowski et al. 2001, 2005), are important for assessing the ploidy of plants, in particular of newly formed hybrid units.

### Size of pollen grains

The lengths of the pollen grains were provided for some species and amphiploids of oat (Katsiotis and Forsberg 1995b; Meo 1999; Chrząstek et al. 2009). Among the accessions studied here, the largest pollen grains were found in an octoploid *A. eriantha* × *A. sativa,* but some hexaploid species differed from values published by Katsiotis and Forsberg (1995b). The differences may be related to the inter-cultivar variation, or the influence of environmental conditions on the size of the pollen grains. The variation of pollen size between seasons can confirm that external factors influenced this characteristic; however, a pattern of inter-OTU differences was maintained (see Table 2, 3).

The size of the pollen is determined by the ploidy level and genotypes of both the gametophyte and the sporophyte in which the pollen grains are produced (Ottaviano and Mulcahy 1989; Southworth and Pfahler 1992; Katsiotis and Forsberg 1995b). Our research on fifteen oat accessions showed a statistically significant correlation between ploidy and length of pollen grains. According to Katsiotis and Forsberg (1995b), the size of pollen grains was determined primarily by the level of ploidy, and only to a small extent by interspecific variation at the same level of ploidy. They proved that there were no significant differences in pollen size between diploid and tetraploid accessions of oats and between hexaploids and octoploids. However, Southworth and Pfahler (1992) found that the size of pollen was influenced not only by the level of ploidy, but also by the number of genomes and volume of cytoplasm, and interactions between these variables. The development of pollen grains in the pollen sac may also be significantly influenced by the orientation of cytokineses in the microspore stem cell and the microspore - tapetum spatial relationship (Kosina and Florek 2011). In grasses, microsporocytes are arranged in one layer which adheres to tapetum (Bhandari 1984; Batygina 1987). Such an architecture of the anther sac provides the equal nutritional support for microspores formed from microsporocytes after two anticlinal cytokineses. Looking from outside into the anther sac, a uniform mosaic of isolateral tetrads of microspores can be seen (Batygina 1987). However, if cytokineses are a combination of anticlinal and periclinal events or there are no cytokineses, so microspores within a tetrad are not equally nourished by tapetum (Brown and Lemmon 2000; Kosina and Florek 2011). Such development creates microspores and pollen grains of various size and mating value. Thus, the large, normal and small microspores form a mosaic at walls of the anther sac. The mosaics were observed in various plant organs (Kosina 2007; Tomaszewska and Kosina 2018), and also in anthers of *Allium senescens* subsp. *montanum* (Małecka 2008). In addition, natural mutations or transgressive segregation may occur in the pollen sac in the genus *Avena,* broadening the old by new variability (Kosina 2015).

### Pollen viability

Pollen grains are considered viable when they are able to germinate on the stigma. There are many different methods for determining pollen viability, and their advantages and disadvantages have been discussed by Dafni and Firmage (2000). The authors suggested the use of several tests simultaneously, but the results obtained by Platje (2003) confirm the effectiveness of simple and quick *in vivo* pollen staining with acetocarmine.

Factors influencing the viability of pollen grains include flower morphology, environmental conditions (temperature, humidity, soil conditions, nitrogen availability, seasonality) and indirect factors (pollen metabolism, number of nuclei, genetic conditions, breeding methods) (Dafni and Firmage 2000). The differences in pollen viability between consecutive growing seasons suggest the influence of external factors on this trait (see Tables 2 and 3). Some amphiploids and parental species studied here showed a reduced viability of pollen grains in one of the growing seasons, which was most likely caused by unfavorable weather conditions, such as high temperature or air humidity. It is known that the early stages of microspore stem cell development are particularly sensitive to high temperature (Sakata et al. 2000). Some of the fifteen examined oat taxa could be particularly sensitive to weather conditions, which resulted in a reduced viability of pollen grains in some growing seasons.

Our studies showed that the percentage of viable pollen was slightly lower in oat amphiploids compared to their parental species. Analysis of pollen viability in *A. fatua* and fatuoids carried out by Warzych (2001) indicated a different percentage of viable pollen between accessions, plants from one accession, and between flowers located in one spikelet. Significant information on the viability of pollen grains was provided by studies on the reproductive system of other grass hybrids. The frequency of viable pollen was significantly lower in hybrids involving *Agrostis* L. and *Polypogon* Desf. compared to the parental species (Zhao et al. 2007). The average pollen viability of interspecific hybrids was higher than that of intergeneric hybrids. A similar tendency was shown by amphiploids resulting from the crossing of *Aegilops ovata* L. with *Secale cereale* L. (Wojciechowska and Pudelska 2002) and *Aegilops kotschyi* Boiss. and *Aegilops biuncialis* Vis. with *S. cereale* (Wojciechowska and Pudelska 2005). However, oat amphiploids appear to be genetically stable and meiotically compatible (Loskutov 2001) despite having distant genomes that exhibit numerous intergenomic translocations (Tomaszewska and Kosina 2018, 2021).The above phenomenon can be explained by the selection shift of the chromosome number, e.g. in the *Avena sativa* × *Avena maroccana* Gand. (*A. maroccana* is a synonym of accepted *Avena magna* H. C. Murphy & Terrell) amphidecaploid (Kushwaha et al. 2004). In this amphidecaploid, the loss of chromosomes over many generations, and simultaneously stable cytotypes with the even number of chromosomes were formed.

### Anomalous pollen development

Reduced viability of pollen grains or even complete sterility, often resulted from anomalous stamen development (Leighty and Sando 1924; Protasevich 1984; Murai and Tsunewaki 1993; Kosina and Florek 2011). No anomalous development of stamens was found in the oat accessions analysed here. Therefore, the observed variability in the viability of pollen grains between the accessions of amphiploids and their parental species may be caused by disturbances in meiosis resulting from the coexistence of several different genomes in the hybrid nuclei (Chen et al. 1977; Tomaszewska and Kosina 2021). Anomalous meiosis can lead to the formation of gametes with a reduced or increased number of chromosomes. Numerous disturbances in the meiotic division, such as delayed chromosomes in telophase I and precocious chromosome segregation, were observed in interspecific and intergeneric hybrids involving *Agrostis* and *Polypogon* (Zhao et al. 2007). The presence of micronuclei and formation of polyads including micropollens were found during microsporogenesis in various young hybrids or amphiploids: *Triticum aestivum* L. × *Agropyron glaucum* Roem. & Schult. (syn.) (Cicin 1978), *Trticum turgidum* L. × *Aegilops squarrosa* L. (synonym of accepted *Aegilops tauschii* Coss) (Kihara 1982), *Triticum aestivum* × *Leymus mollis* (Trin.) Pilg. (Li et al. 2005). Disturbances in meiosis were also observed in interspecific oat hybrids (Paczos-Grzęda 2003) and some oat species, including diploids (Sheidai et al. 2003). These include chromosomes delayed in anaphase I and II and telophase I and II, micronuclei present in the microspore tetrads, bridges, chromosome elimination, multipolar systems and cytomixis. Examples of similar disorders came from the meiotic analysis of the *Zea mays* L. inbred lines (Caetano-Pereira and Pagliarini 2001) in which premature chromosome segregation, delayed chromosomes and micronuclei have been observed. The participation of such disturbed gametes in fertilization may contribute to the formation of an aneuploid generation with reduced fertility and viability (Zhao et al. 2007).

Numerous examples of anomalous pollen grains can be found in the literature. Micropollens and ultra-micropollens seem to be quite common in oats (Kosina and Florek 2011), and may arise through asymmetric segregation of chromosomes (Baptista-Giacomelli et al. 2000). Irregular pollen grains can be formed by abnormal pollen wall formation, unfinished cytokinesis, and cytomixis. Similar disorders were observed in an amphiploid *Triticum aestivum* × *Leymus mollis* (Li et al. 2005). Many cytomixis cases were found in *Diplotaxis harra* (Forssk.) Bois., where aneuploid cells showed highly unequal number of chromosomes (Malallah and Attia 2003). Abnormal cell wall formation is one of the most common causes of large, non-reduced microspores (Ellison 1937). Anomalous cytokinesis may occur during mitosis or meiotic division I and II. In the hybrid *A. barbata* × *A. strigosa* subsp. *hirtula* (Lag.) Malzev (syn. of *Avena strigosa* Schreb), pollen binuclear stem cells were observed, the presence of which resulted from disturbances in cell wall formation during pre-meiotic mitosis (Holden and Mota 1956). The second nucleus, usually peripheral, underwent apoptosis preceded by chromatin fragmentation and nucleolus atrophy. Diploid gametes have also been observed in other species and hybrids of the genus *Avena* (Ellison 1937; Kosina and Florek 2011), and diploid gametes accounted for 1% of the pollen grains in tetraploid *Avena vaviloviana* Malzev (Mordv.) (Katsiotis and Forsberg 1995a). Large, unreduced microspores can be formed by cytomixis or restitution of nuclei during meiosis (Stuczyński et al. 1994). Unreduced pollen grains have low mating value and can be very ineffective in the fertilization process due to slow growth of their thick pollen tubes, as it was evidenced in *Prunus spinosa* L. (Staszak 2004).

### Number of poruses

Morphogenesis of poruses seem to be related to *de novo* synthesis of callose in a microspore (Teng et al. 2005). In grasses, a pattern of callose deposition in a differentiating pollen grain appeared to be different. Scattered lenses of callose adjacent to intine were observed in *Brachypodium phoenicoides* (L.) Roem. & Schult. (Kłyk 2005), while in *Avena fatua* callose was synthesised around a pore and disappeared towards the opposite pole (Warzych 2001).

The number of poruses in the pollen wall, through which pollen tubes can germinate, varies between species. Pollens with different number of apertures often occur in the same species, including *Viola diversifolia* (DC.) W. Becker (Dajoz et al. 1991), *Viola arvensis* Murray and *Nicotiana tabacum* L. (Ressayre et al. 1998). The grass group is characterised by single-porate pollen grains, thus pollen with an increased number of poruses should be considered anomalous. Multiporate pollen grains were observed in all amphiploids studied here, while their parental species produced pollen with only one porus. Multiporate pollen grains were found in different hybrids and amphiploids of grasses: diporate grains were developed in an F1 hybrid generation of *Triticum turgidum* × *Aegilops squarrosa* (Kihara 1982), half of the pollen pool of *Miscanthus* Anderss. ‘Giganteus’ had two to five poruses (Linde-Laursen 1993), pollen with ten poruses has been reported in an amphiploid *Triticum aestivum* × *Leymus mollis* (Li et al. 2005). It should be emphasized that multiporate pollen grains can germinate through many pores. As a result, competition occurs between the tubes during their growth, which significantly reduces their effectiveness in the fertilization process. Such a case was observed in the amphiploid *Avena barbata* × *A. sativa* subsp. *nuda* (Florek 2013; Kosina et al. 2014).

The increase number of poruses may be caused by the independent activity of genes located in different genomes (Kalinowski et al. 2001, 2005). Therefore, plants formed by polyploidisation are often characterised by multiporate pollen grains. Some authors discuss the relationship between apomictic mode of reproduction and multiporate pollen. Liu et al. (2004) recorded multiporate pollen grains in *Leptochloa panicea* (Retz.) Ohwi and *Schmidtia pappophoroides* Steud. ex J.A.Schmidt, which showed apomictic type of reproduction. Other apomicts among grasses that produce multiporate pollen include genera *Apluda* L., *Heteropogon* Pers., *Panicum* L., *Paspalum* L., *Pennisetum* Rich. (Guohua et al. 2009) and *Bothriochloa* Kuntze (Ma and Huang 2007). All of these genera include species showing high ploidy levels (Higgins et al. 2021; Tomaszewska et al. 2021a,b). It can be concluded that formation of multiporate pollen is associated with young allopolyploids and the interaction of various genomes in the nucleus, whereas apomictic reproduction allows the population to function with a generative defect. Allopolyploids often showed numerous disturbances in micro- and macrosporogenesis, which can lead to apomixis. In *Bothriochloa ischaemum* (L.) Keng, cytogenetic analyses revealed disturbed chromosome segregation and delayed chromosomes (Ma and Huang 2007). In *Calamagrostis hakonensis* Franch. & Sav. and *Dichanthium aristatum* (Poir.) C.E.Hubb (Guohua et al. 2009), double fertilisation was impossible due to sterility of pollen, thus apomixis was an alternative to sexual reproduction. It was found that a shift from the generative reproduction to apomixis occurred in plant populations showing higher numbers of chromosomes, where meiotic disorders and defective pollens appeared (Grant 1981). Thus, the interaction between the development of multiporate pollen grains and apomictic reproduction requires further research to understand what is the cause and what is the effect.

## Concluding remarks

The pollen of the examined oat species and the amphiploids obtained from their crosses showed high viability, which can result in a proper development of caryopses. However, the pollen viability of amphiploids was several percent lower than in species. In addition, anomalous pollens were produced at low percent by the amphiploid flowers. It was found that the size of pollen grains varied during the growing seasons, but the interrelations between the studied taxa were stable. The viability of pollen grains was more significantly affected by the environment than the length of pollen grain and showed no correlation with the variability of the grain size. In the ordination space, amphiploids were discriminated from parental species. In both groups of plants, a positive correlation was maintained between the pollen size and the level of ploidy, however, in the cloud of correlation points along the regression line, amphiploids were located among species with a high level of ploidy and were extreme units there. Developmental anomalies, shown in pollen morphotypes, had a low frequency, among them the formation of pollen micrograins was most common. Such an event proved that some pollen grains were chromosomally unbalanced and were formed in polyads. Anomalies were more common in the hybrid types, and the formation of pollen grains with many poruses occurred only in amphiploids and was strongly correlated with the frequency of micropollens. Multiporate grains will be less efficient for reproductive success due to their germination with many pollen tubes and mutual competition (Florek 2013; Kosina et al. 2014). In the ordination space, monoporate types (species) were well discriminated from multiporate types (amphiploids). The high level of pollen viability in amphiploids proved their probable genomic/chromosomal stabilisation through many generations of their reproduction in the life collections.

## Declarations

### Funding

Statutory funds (1232/M/IBR/11) supported by University of Wrocław, Poland, and support from the European Union’s Horizon 2020 research and innovation programme under the Marie Sklodowska-Curie grant agreements No 844564 and No 101006417 for PT.

**Compliance with ethical standards**

### Conflict of interest

The authors declare that they have no conflict of interest.

### Ethics approval

The authors guarantee compliance with ethical standards.

### Consent to participate

Not applicable

### Consent for publication

Not applicable

### Availability of data and material

Not applicable

### Code availability

Not applicable

### Abbreviations

A file included

## Authors’ contributions

PT designed and conducted all analyses and interpreted the results. PT and RK prepared figures and wrote the article. RK provided research idea, supervised the experiments and research documentation. The authors read and approved the manuscript.

## Acknowledgments

We are grateful to Bundesanstalt für Züchtungsforschung an Kulturpflanzen, Braunschweig, Germany; National Small Grains Collection, Aberdeen, Idaho, USA; and Vavilov Institute of Plant Industry, St. Petersburg, Russia for their generous provision of seeds.

## Notes

### Competing Interest Statement

The authors have declared no competing interest.

